# En1 and Lmx1b do not recapitulate embryonic dorsal-ventral limb patterning functions during mouse digit tip regeneration

**DOI:** 10.1101/2022.03.15.484493

**Authors:** Gemma L. Johnson, Morgan B. Glasser, Julia F. Charles, Jeffrey Duryea, Jessica A. Lehoczky

**Affiliations:** Department of Orthopedic Surgery, Brigham and Women’s Hospital, Boston, MA, 02115, USA; Department of Systems Biology, Harvard Medical School, Boston, MA, 02115, USA; Department of Medicine, Brigham and Women’s Hospital, Boston, MA, 02115, USA; Department of Radiology, Brigham and Women’s Hospital, Boston, MA, 02115, USA

**Keywords:** Digit tip regeneration, dorsoventral patterning, limb development, En1, Lmx1b

## Abstract

The mouse digit tip is a complex tissue that is capable of regeneration after amputation. How the regenerated digit tip is patterned is unknown, but a long-standing hypothesis in the field of regeneration proposes that developmental patterning mechanisms are re-used during regeneration. The digit tip bone exhibits strong dorsal-ventral (DV) polarity, so we focus on Engrailed 1 (En1) and LIM homeobox transcription factor 1B (Lmx1b), two well-studied transcription factors necessary for DV patterning during limb development. We investigate if En1 and Lmx1b are re-expressed during regeneration in a developmental-like spatially restricted pattern, and if they direct DV morphology of the regenerated digit tip. We find that both En1 and Lmx1b are expressed in the regenerating mouse digit tip epithelium and mesenchyme, respectively, but without DV polarity. We use conditional genetics and quantitative analysis of digit tip bone morphology to determine that genetic deletion of En1 or Lmx1b in adult digit tip regeneration modestly reduces bone regeneration but does not affect DV patterning of the regenerate. Collectively, our data suggest that while En1 and Lmx1b are re-expressed during mouse digit tip regeneration, they do not define the DV axis during regeneration.

## INTRODUCTION

Some tetrapods, including salamanders, newts, and juvenile xenopus, are able to regenerate entire limbs after amputation, and several species of fish, exemplified by zebrafish, are capable of fin regeneration (Johnson and Weston, 1995; Spallanzani, 1768). Humans and mice do not regenerate limbs but do innately regenerate amputated digit tips, the most distal structures of the limb (Borgens, 1982; Illingworth, 1974; Zhao and Neufeld, 1995). Limb and appendage regeneration in all of these models requires the blastema, a collection of proliferating progenitor cells, which forms underneath the wound epithelium covering the amputation site (Carlson, 1978; Hay and Fischman, 1961; Morgan, 1901). This blastema grows and differentiates into a new limb or appendage which re-establishes the correct three-dimensional pattern and tissue organization of the original structure in the proximal-distal (PD), anterior-posterior (AP), and dorsal-ventral (DV) axes. However, despite extensive study in salamanders, it remains unclear how the correct patterning is restored in the regenerating limb (reviewed in (Flowers and Crews, 2020; Vieira and McCusker, 2019)).

Embryonic limb development is a well-studied model of how multipotent mesenchyme differentiates into the tissues of the mature limb in a stereotyped three-dimensional pattern. Because there are strong similarities in terms of tissue types and their interactions, the process of limb regeneration is often compared to embryonic limb development, which provides a useful lens for studying morphogenesis during regeneration (Goss, 1969). The developing limb bud is comprised of undifferentiated mesenchymal cells surrounded by the apical ectodermal ridge (AER), a specialized distal epithelium that is integral for the proliferation and differentiation of the underlying mesenchyme (Saunders, 1948; Summerbell, 1974). As with the developing limb bud, the regenerative blastema in salamander, frog, and mouse is largely made up of mesenchymal cells that are surrounded by a specialized distal epithelium, the wound epithelium, that signals to the underlying mesenchyme (Aztekin et al., 2021; Boilly and Albert, 1990; Campbell and Crews, 2008; Endo et al., 2000; Ghosh et al., 2008; Lee et al., 2013; Satoh et al., 2008; Storer et al., 2020). Furthermore, in both limb development and regeneration, progenitor cells proliferate and re-pattern to form a structure that is the correct size and shape in comparison to the rest of the organism. Beyond tissue types and morphology, there is also evidence for shared gene expression patterns between limb regeneration and development. In the axolotl, posterior HoxA genes are re-expressed during limb regeneration in the same spatial and temporal pattern as during limb development (Gardiner et al., 1995; Roensch et al., 2013). In mice, Fgf2 is expressed in the AER as well as in the wound epithelium of regenerating digit tips (Takeo et al., 2013). In addition, the cell types involved in limb development and regeneration have been compared by single-cell RNA-sequencing analyses in several model systems (Aztekin et al., 2021; Gerber et al., 2018; Storer et al., 2020). In mice, for example, a subset of mesenchymal digit tip blastema cells do express embryonic limb development related genes such as Hoxa13 and Hoxd13, as well as many genes related to ossification (Qu et al., 2020; Storer et al., 2020), but their overall transcriptional profiles are not equivalent to those of embryonic limb bud cells (Storer et al., 2020). The mouse wound epithelium has not been directly compared to the AER. However, the specialized wound epithelium during frog regeneration is transcriptionally similar to the AER (Aztekin et al., 2021). While these studies show developmental limb patterning genes are re-expressed during regeneration, it remains unknown whether they have the same functional role in regeneration as they do in development.

Mouse digit tip regeneration is a robust and human-relevant model to dissect the role of limb development genes during regeneration at a genetic and functional level. The digit tip is a distal limb structure comprised of several different tissue types including bone, connective tissue, blood vessels, nerves, epithelium, and the nail. Amputating the distal mouse digit tip results in a stereotyped regenerative process which includes formation of a blastema made up of several different cell types including a heterogeneous population of fibroblasts, pre-osteoblasts, Schwann cells, immune cells, and vascular smooth muscle and endothelial cells (Fernando et al., 2011; Johnson et al., 2020; Marrero et al., 2017; Simkin et al., 2017; Storer et al., 2020). While the digit tip blastema is not identical in cell type to mouse embryonic limb bud cells (Storer et al., 2020), it remains to be determined if blastema cells are under the influence of limb development molecular pathways in the patterning of the regenerating digit tip. For example, genes necessary for embryonic limb patterning may be re-deployed in morphogenesis of the regenerating mouse digit tip.

In this paper, we expand the comparison of mouse digit tip regeneration to limb development through the lens of patterning. Of the PD, AP, and DV axes, the digit tip bone has the most phenotypic variation in the DV axis. We exploit the distinct DV bone morphology to investigate the role of Engrailed 1 (En1) and Lim homeobox transcription factor 1 beta (Lmx1b), two transcription factors necessary for DV patterning in the developing limb bud, during digit tip regeneration. Using RNA in situ hybridization, we define the expression domains of En1 and Lmx1b during digit development and regeneration, showing that En1 and Lmx1b are expressed during mouse digit tip regeneration but with no DV restriction. Conditional genetics and a custom computational analysis of microCT scans of 490 digit tip bones enable us to determine that loss of En1 or Lmx1b modestly impairs bone regeneration but does not alter the DV morphology of regenerated digit tip bones. Taken together, our data show that En1 and Lmx1b do not direct DV patterning during digit tip regeneration, challenging a long-standing hypothesis that regeneration re-deploys developmental patterning genes (Goss, 1969).

## RESULTS

### Dorsally restricted Lmx1b expression persists throughout digit development while En1 expression is dynamic

The expression domains of En1 and Lmx1b in early mouse limb development are well described. Lmx1b expression is restricted to the dorsal limb bud mesenchyme by embryonic day (e) 10.5 (Cygan et al., 1997; Loomis et al., 1998) (Sup Fig 1C, C’). This expression persists in the dorsal mesenchyme of the distal autopod, including the digit, until at least e16.5 (Dreyer et al., 2004). Whether Lmx1b expression in the digit persists after e16.5, and in which tissues, is not reported. Separately, it has been established that En1 is expressed in the ventral limb bud epithelium by e9.5, as soon as the limb bud is formed (Davis et al., 1991) (Sup Fig 1B, B’). En1 continues to be expressed in the digit epithelium until at least e13.5 (Loomis et al., 1998), but the expression pattern of En1 in the digit after this stage is not reported. To determine if expression of En1 and Lmx1b persists in the developing digit tip and if they maintain embryonic limb bud DV restriction, we used hybridization chain reaction RNA fluorescent in situ hybridization (HCR RNA-FISH) (Choi et al., 2018) with probes to Lmx1b and En1 in e16.5 (embryonic), postnatal day 4/5 (PN4/5; neonatal), and 6 week old (adult) digits (Fig 1A). HCR RNA-FISH in the e16.5 digit shows that En1 is expressed in the ventral epithelium (Fig 1D, D’) with some expression dorsal to the DV boundary at the distal tip of the digit (Fig 1C, C’), but not in the rest of the dorsal epithelium (Fig 1B, B’). By PN4/5, epithelial En1 expression is maintained ventrally (Fig 1G, G’) and at the distal tip (Fig 1F, F’) and has expanded further dorsally (Fig 1E, E’) but does not yet reach the nail matrix, a more proximal region of the dorsal epithelium (not shown). In the adult digit, no DV En1 restriction is found, and expression is detected throughout the entirety of the digit tip epithelium (Fig 1H’-J’). Thus, En1 is expressed exclusively in the epithelium, and the original ventral expression domain expands dorsally as the digit matures.

**Figure 1.**
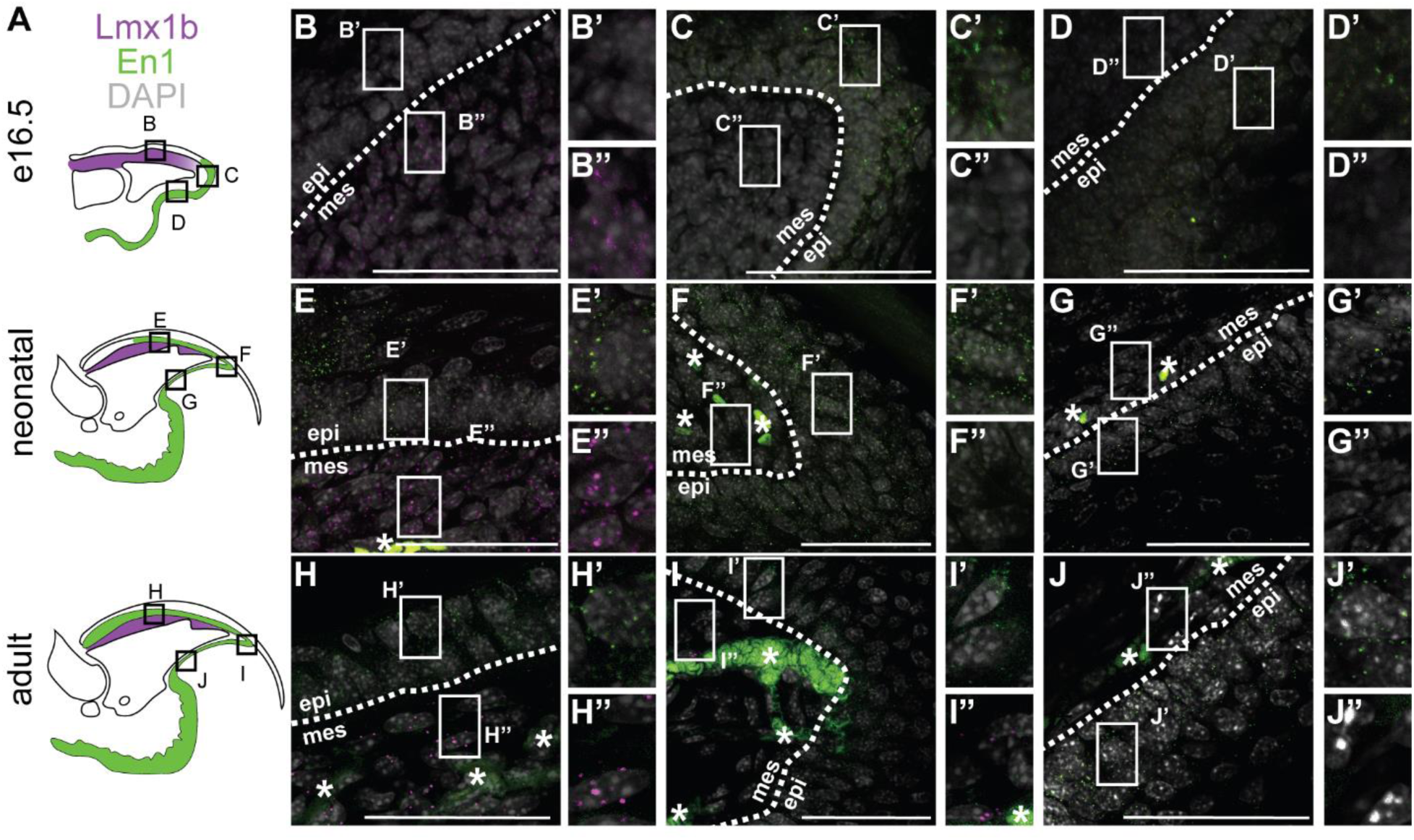
Expression of En1 and Lmx1b during digit tip development. (A) Schematics of mouse digit tip cross sections at e16.5 (embryonic; top), P4/5 (neonatal; middle) and 8 weeks old (adult; bottom), including En1 expression in green and Lmx1b expression in magenta. Lettered boxes correlate with position of in situ images on right. (B-J) HCR RNA-FISH for En1 (green puncta) and Lmx1b (magenta puncta) in embryonic (B-D), neonatal (E-G), and adult (H-J) digit tip sections. (A-C). DAPI is shown in grey. Scale bars represent 50μm. Asterisks (*) denote blood vessel and red blood cell autofluorescence. Dashed lines show epithelial border. White boxes show position of magnified panels to the right. Epi = epithelium, mes = mesenchyme.

Conversely, we find that Lmx1b is expressed in the dorsal (Fig 1B, B’’) but not distal (Fig 1C, C’’) or ventral (Fig 1D, D’’) mesenchyme in the e16.5 digit. In the neonatal digit, Lmx1b continues to be expressed in dorsal mesenchyme (Fig 1E, E’’) but not at the distal tip of the digit (Fig 1F, F’’) or ventrally (Fig 1G, G’’). Similarly, in the adult digit tip, Lmx1b is expressed in the dorsal mesenchyme (Fig 1H, H’’) and has extended more distally to the tip (Fig 1I, I’’) but is not expressed in the ventral mesenchyme (Fig 1J, J’’). At all of these developmental stages, Lmx1b is expressed only in mesenchymal cells and not in the epithelium (Fig 1A, B’-J’).

During mouse limb development, En1 and Lmx1b are part of a network of gene expression that includes Wnt7a (Chen and Johnson, 2002; Loomis et al., 1996; Parr and Mc Mahon, 1995; Parr et al., 1993). En1 is expressed in the ventral epithelium and decreases expression of Wnt7a, leading to Wnt7a expression only in the dorsal epithelium. Wnt7a then induces Lmx1b expression in the adjacent dorsal mesenchyme (Sup fig 1A). Wnt7a was detected in the epithelium of control e12.5 forebrain sections by HCR RNA-FISH (Sup Fig 3A, A’, B) but HCR RNA-FISH of adult digits for Wnt7a shows no expression in the epithelium or otherwise in the digit (Sup fig 3C-G). Taken together our data show that expression of both En1 and Lmx1b persists throughout digit development and into adult digit homeostasis, whereas Wnt7a expression is undetectable in the adult. Despite the change in the En1 expression domain throughout development of the digit tip, En1 remains restricted to the epithelium, as in early limb bud development, and Lmx1b remains restricted to the mesenchyme.

### En1 and Lmx1b are expressed in the regenerating digit tip without dorsal-ventral polarity

En1 and Lmx1b expression persists into adulthood in the digit tip in the same tissue compartments as during limb development (Fig 1), suggesting that they could function in DV patterning during digit tip regeneration. To address this, we first turned to our single-cell RNA-sequencing (scRNA-seq) data from adult mouse regenerating digit tips (Johnson et al. 2020, accession GSE143888) to determine if En1 and Lmx1b are expressed during digit tip regeneration. Combining scRNA-seq data from 11, 12, 14, and 17 days postamputation (dpa) and unamputated controls, 95.4% of cells expressing En1 are epithelial and 95.1% of cells expressing Lmx1b are fibroblasts (98.7% are of general mesenchymal origin) (Fig 2A, 2B), consistent with their tissue-specificity during limb development. To assess any spatial restriction of En1 and Lmx1b expression during regeneration and whether it recapitulates that of early limb development, we utilized HCR RNA-FISH for Lmx1b and En1 in the regenerating digit tip at 11dpa (Fig 2D-I). 11dpa represents an early stage of regeneration when the blastema has formed but no there is no widespread differentiation (Fig 2C middle) (Fernando et al., 2011). Consistent with the scRNA-seq data, HCR RNA-FISH shows that En1 is expressed contiguously throughout the dorsal and ventral epithelium of the regenerating digit tip (Fig 2D, D’, F, F’). En1 expression is also present in the epithelium at the distal tip surrounding the blastema (Fig 2E, E’), and thus deviates from the DV spatial restriction found in the embryonic limb bud. This suggests that the function of En1 in regeneration could be discrete from that of limb development.

**Figure 2.**
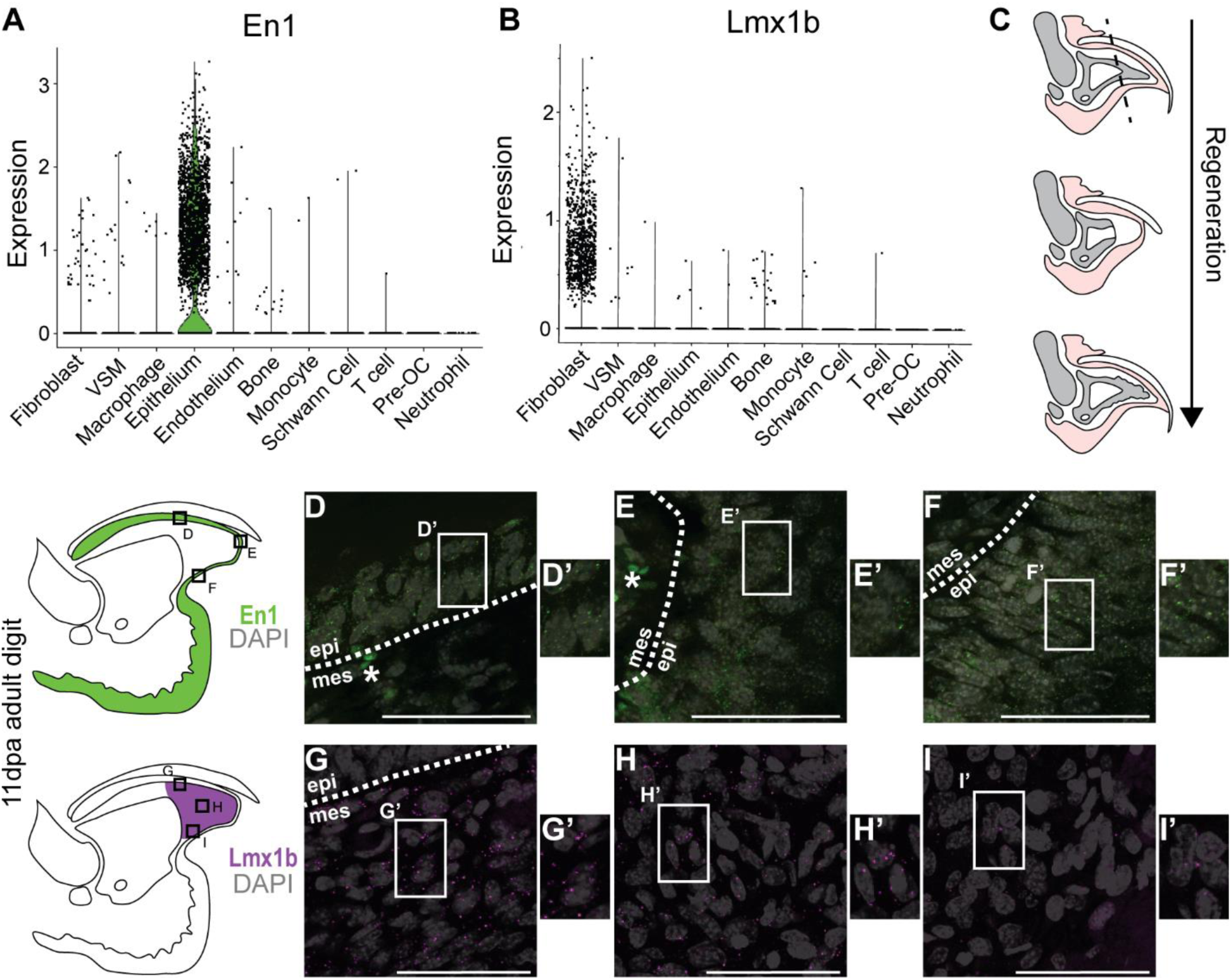
Expression of En1 and Lmx1b during digit tip regeneration. (A-B) Gene expression of En1 (A) and Lmx1b (B) in regenerating and unamputated digit tips shown by violin plot. Black dots represent individual cells. (C) Schematic of mouse digit tip regeneration. Digits are amputated at the plane of the dashed line (top), resulting in blastema formation (middle) and regeneration of distal digit tip structures (bottom). (D-I) HCR RNA-FISH for En1 (green puncta) (D-F) and Lmx1b (magenta puncta) (G-I) in 11dpa regenerating digit tips. Schematics of mouse digit tip cross sections at 11dpa are to the left and include En1 expression in green and blastema specific Lmx1b expression in magenta. Lettered boxes correlate with position of in situ images on right. White boxes show location of magnified inset to the right. DAPI is shown in grey. Scale bars represent 50μm. Asterisks (*) denote blood vessel and red blood cell autofluorescence. Dashed lines show epithelial border. Epi = epithelium, mes = mesenchyme.

By HCR RNA-FISH, Lmx1b is heterogeneously expressed at a low level in cells throughout the blastema at 11dpa and not found in the epithelium (Fig 2G-I). Lmx1b is expressed by cells in the dorsal blastema (Fig 2G, G’) as well as the central (Fig 2H, H’) and ventral (Fig 2I, I’) regions of the blastema. This pattern holds through later stages of digit tip regeneration: Lmx1b continues to be expressed dorsally (Sup Fig 2A, A’, D, D’), centrally (Sup Fig 2B, B’, E, E’), and ventrally (Sup Fig 2C, C’, F, F’) throughout the mesenchyme of the blastema at both 14dpa (Sup Fig 2A-C) and 17dpa (Sup Fig 2D-F). This is in contrast to embryonic limb bud and adult homeostatic digit Lmx1b expression which is restricted to dorsal mesenchyme, suggesting a regeneration-specific role for Lmx1b. However, Lmx1b is expressed in the mesenchyme, a tissue type consistent with limb bud development and the scRNA-seq data. In contrast to early limb development (Sup Fig 1A), Wnt7a expression was not detected in the epithelium of either unamputated (UA) or 11dpa regenerating digit tips (Sup Fig 3H-K), confirming the lack of Wnt7a expression found in the scRNA-seq data from combined epithelium of UA and regenerating digit tips (Johnson et al., 2020) (Sup Fig 3C). These expression studies suggest that the embryonic limb DV patterning network is not re-deployed during digit tip regeneration to direct DV morphogenesis.

### Embryonic loss of En1 and Lmx1b results in dysmorphic digits and affects digit tip regeneration

To functionally test if En1 or Lmx1b are necessary for DV patterning during digit tip regeneration, we utilized tissue specific conditional genetics. We first bred En1-flox;K14-cre mice, which have genetic deletion of En1 in the epithelium (Dassule et al., 2000; Sgaier et al., 2007). As previously reported, embryonic loss of En1 in the epithelium results in limbs absent of ventral structures and with duplicated dorsal structures such as hair follicles on the palm and nails extending to the ventral digit (Loomis et al., 1996) (Fig 3A,D). Consistent with this phenotype, En1-fl/fl;K14-cre digit tip bones are cylindrical, exhibiting “double dorsal” morphology and a loss of ventral structures such as the sesamoid bone and ventral hole (Fig 3E), as compared to En1-wt/wt;K14-cre digits (Fig 3B). To determine the effect of developmental loss of En1 on digit tip regeneration, we amputated these digits at PN3 and evaluated regenerated digit tip bones at 3 weeks post amputation (wpa). We calculated the percent regeneration by dividing regenerated digit length by the same animal’s contralateral UA digit length. Both En1-fl/fl;K14-cre and En1-wt/wt;K14-cre digit tip bones regenerated following amputation (Fig 3C,F). Surprisingly, the En1-fl/fl;K14-cre regenerated digits had longer bones than the wildtype controls (Fig 3M) and appear to regenerate with double dorsal morphology (Fig 3F). However, because the digit tip bone was dysmorphic in the DV axis prior to amputation, it is unclear whether the aberrant regenerated bone morphology is due to loss of En1 or due to the pre-existing morphology of the bone and surrounding tissues.

**Figure 3.**
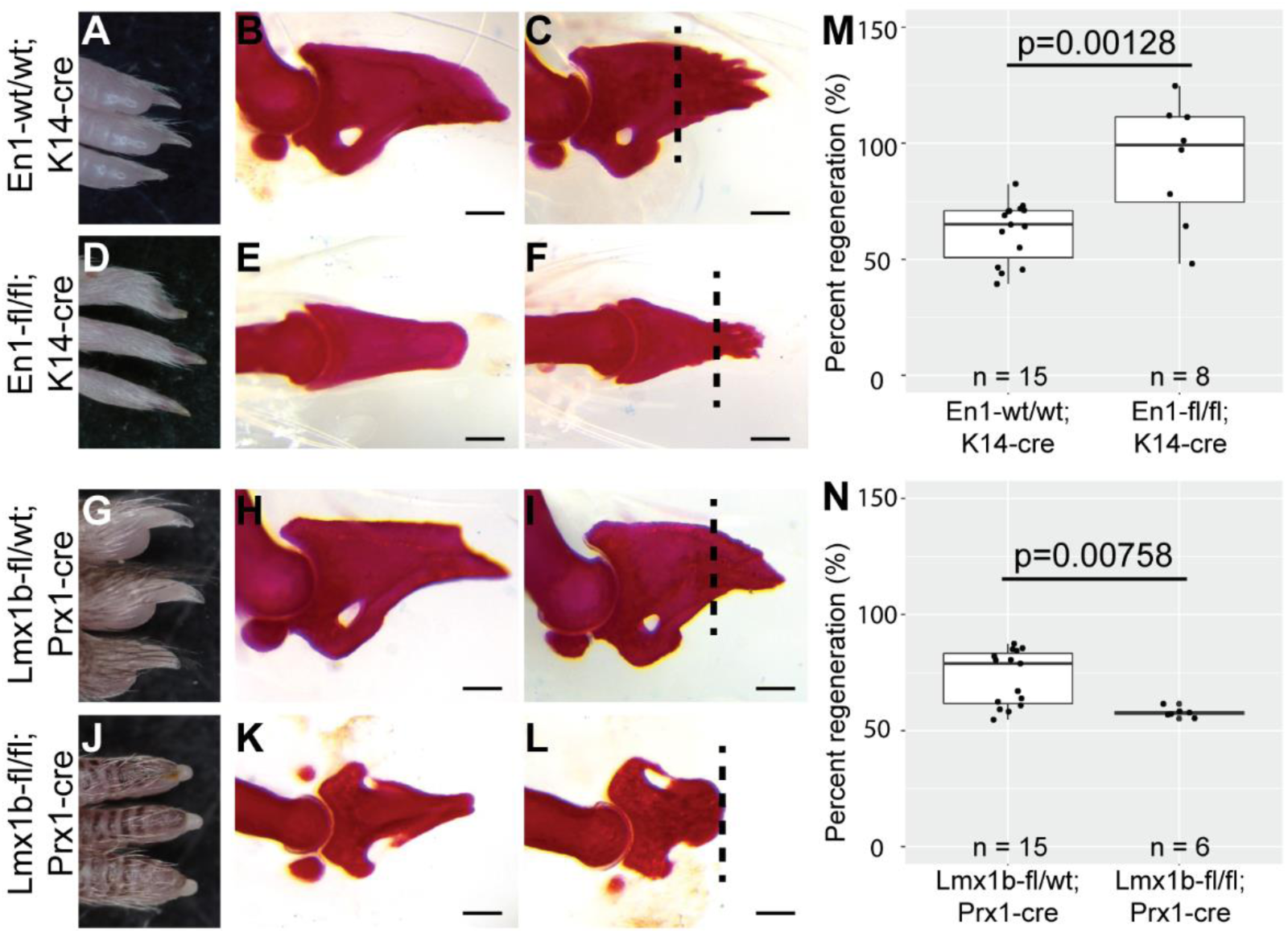
Developmental loss of En1 or Lmx1b perturbs digit tip regeneration. (A-C) Gross morphology ventral view (A) of control En1-wt/wt;K14-cre digits and alizarin red stained unamputated (B) and 3 wpa (C) En1-wt/wt;K14-cre digit bones. (D-F) Gross morphology ventral view (D) of En1-fl/fl;K14-cre digits and alizarin red stained unamputated (E) and 3 wpa (F) En1-fl/fl;K14-cre digit bones. (G-I) Gross morphology dorsal view (G) of control Lmx1b-fl/wt;Prx1-cre digits and alizarin red stained unamputated (H) and 3wpa (I) Lmx1b-fl/wt;Prx1-cre digit bones. (J-L) Gross morphology dorsal view (J) of Lmx1b-fl/fl;Prx1-cre digits and alizarin red stained unamputated (K) and 3wpa (L) Lmx1b-fl/fl;Prx1-cre digit tip bones. Dashed lines denote amputation plane. Scale bars represent 200um. (M-N) Box plot of digit tip bone percent regeneration in En1-wt/wt;K14-cre or En1-fl/fl;K14-cre 3wpa digits (M) or Lmx1b-fl/wt;Prx1-cre or Lmx1b-fl/fl;Prx1-cre 3wpa digits (N). n represents number of individual digits shown. Lengths are normalized to contralateral unamputated control digits. * p < 0.05, ** p < 0.01, *** p < 0.001.

We next bred Lmx1b-flox;Prx1-cre mice, which have genetic deletion of Lmx1b in the developing limb (Logan et al., 2002; Zhao et al., 2006). Consistent with previous reports, genetic deletion of Lmx1b in the limb results in aberrant DV limb morphology (Chen et al., 1998). Compared to control Lmx1b-fl/wt;Prx1-cre digits (Fig 3G), Lmx1b knockout mice exhibit loss of dorsal structures such as the nail and gain of ectopic ventral structures like the toe pad on the dorsal side of the digit (Fig 3J). Similarly, the digit tip bones show a “double ventral” phenotype exemplified by an ectopic sesamoid bone and bone hole on the dorsal side (Fig 3K). To test the necessity of Lmx1b for DV morphology in digit tip regeneration we amputated the digit tips of Lmx1b-fl/fl;Prx1-cre PN3 mice and Lmx1b-fl/wt;Prx1-cre controls. At 3wkpa, Lmx1b-fl/wt;Prx1-cre control digit bones regenerate normally (Fig 3I), whereas Lmx1b-fl/fl;Prx1-cre digit tip bones do not regenerate (Fig 3L and 3N). This suggests that Lmx1b is necessary in the mesenchyme for digit tip regeneration. However, Lmx1b-fl/fl;Prx1-cre mice have reduced or absent nails (Fig 3J), a dorsal structure that is necessary for regeneration (Mohammad et al., 1999; Zhao and Neufeld, 1995), so the genetic necessity of Lmx1b for regeneration in this experiment is confounded.

### Conditional deletion of En1 and Lmx1b during regeneration causes modest bone loss

To disentangle the confounding developmental digit tip bone morphologies from any regenerative phenotype, we conditionally deleted En1 and Lmx1b in adult mice with normally developed limbs. We first bred En1 epithelial-specific conditional knockout (cKO) mice using an En1-floxed allele (Sgaier et al., 2007) and a K14-creERT2 tamoxifen inducible cre allele (Li et al., 2000) (Fig 4A). To test the role of En1 during adult digit tip regeneration, we induced recombination in En1-fl/fl;K14-creERT2 mice with three daily doses of tamoxifen and compared them to untreated En1-fl/fl and En1-fl/fl;K14-creERT2 control cohorts. Tamoxifen dosing was sufficient for recombination of the En1 locus, although ectopic recombination was detected in uninduced En1-fl/fl;K14-creERT2 mice (Sup Fig 4E). This genotypic finding is consistent with the uninduced En1-fl/fl;K14-creERT2 phenotype, whereby around 5 weeks of age En1-fl/fl;K14-creERT2 mice, regardless of tamoxifen induction, develop ectopic ventral nails from their toe pads (Sup Fig 4A-B). Despite the leaky cre affecting ectodermal structures in the digit tip, the bone structure is still comparable to En1-fl/fl mice with no evidence of aberrant DV patterning (Fig 4B, 4D, Sup Fig 4C-D). Digit tips of 4 week old mice were amputated one day after the last tamoxifen dose and collected at 4wpa, when adult digit tip bone regeneration is complete, and scanned by micro-computed tomography (uCT) to evaluate bone regeneration (Fig 4A) (Dawson et al., 2018; Fernando et al., 2011).

**Figure 4.**
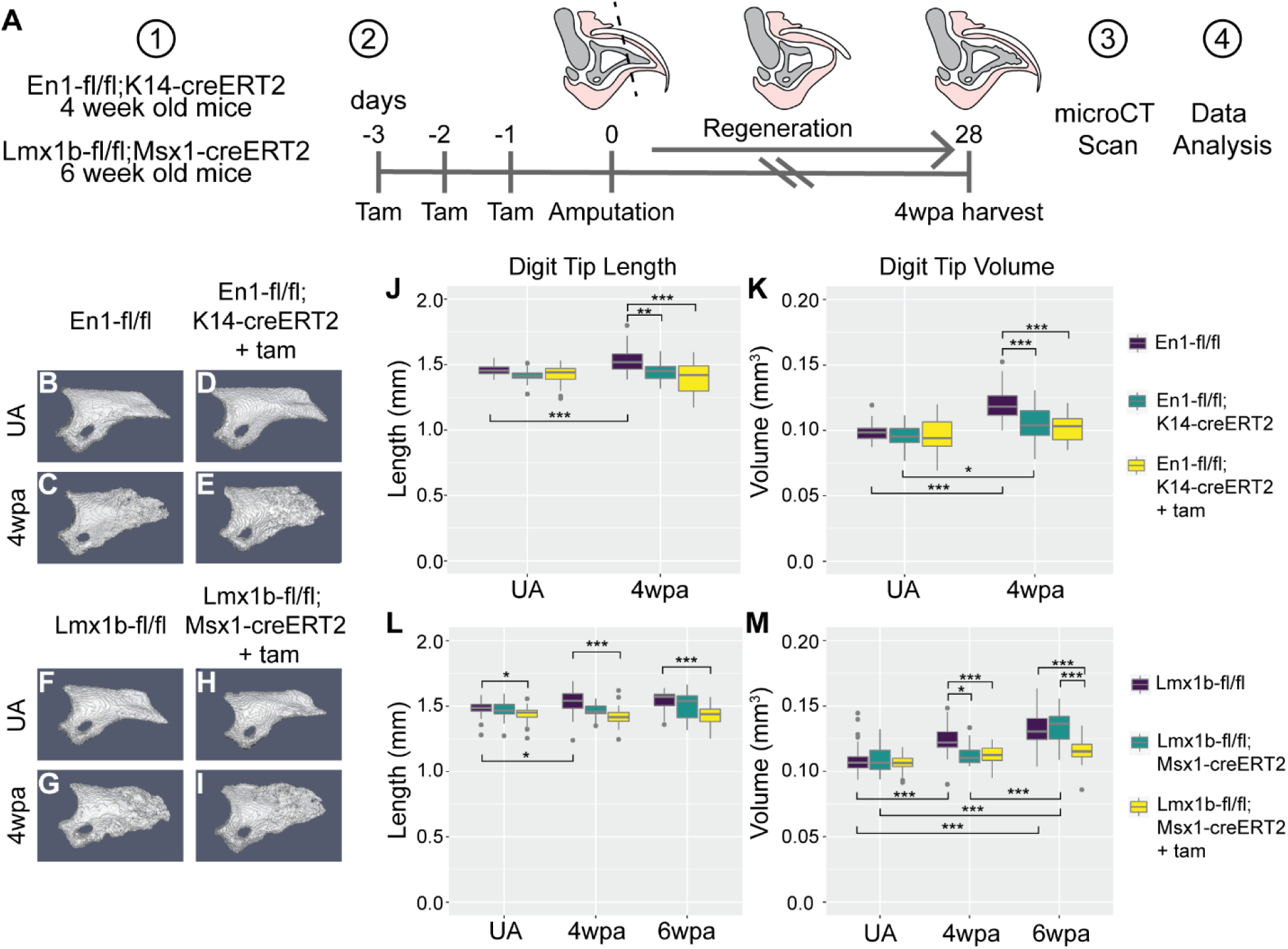
Conditional knockout of En1 and Lmx1b has a small effect on bone regeneration. (A)) Schematic of the En1 and Lmx1b conditional knockout experiment. Experimental cohorts at 4 weeks old (En1-fl;K14-CreERT2) and 6 weeks old (Lmx1b-fl;Msx1-CreERT2) were given tamoxifen by oral gavage for three days, then digits were amputated on the next day. Digits were collected at 4wpa when bone regeneration is considered finished. Digits were scanned by uCT and the resulting data analyzed in a custom software pipeline. (B-I) Representative 3D models of uCT scanned digits. (B-C) En1-fl/fl control UA (B) and 4wpa (C) digits. (D-E) En1-fl/fl; K14-CreERT2 cKO UA (D) and 4wpa (E) digits. (F-G) Lmx1b -fl/fl control UA (F) and 4wpa (G) digits. (H-I) Lmx1b -fl/fl;Msx1-CreERT2 cKO UA (H) and 4wpa (I) digits. (J-M) Box plots showing quantification of digit length and volume for En1 control and cKO digits (J-K) and Lmx1b control and cKO digits (L-M). Number of individual digit tip bones in each group is noted in methods and supplemental figure 5. * p < 0.05, ** p < 0.01, *** p < 0.001.

By 4wpa, both En1-fl/fl;K14-creERT2 induced and En1-fl/fl control digit tip bones successfully regenerated, as grossly assessed from uCT renderings (Fig 4C, 4E). Quantification of length and volume provided more detailed metrics of the regenerated bones. In agreement with previous analyses of wild type digit tip bone length and volume after regeneration by uCT (Dawson et al., 2018; Fernando et al., 2011), the regenerated En1-fl/fl control digits have an over-regeneration of bone length (p = 1.5×10^-4^) and volume (p < 1×10-7) compared to the UA contralateral controls (Fig 4J-4K). En1-fl/fl;K14-creERT2 cKO 4wpa digits have a reduction in length (p < 1×10-7) and volume (p < 1×10-7) compared to regenerated En1-fl/fl control digits (Fig 4J-K). Uninduced En1-fl/fl;K14-creERT2 digits also exhibit reduced regenerative length (p = 0.001) and volume (p = 2.5×10^-6^) compared to En1-fl/fl control digits, consistent with the ectopic recombination found in these digits (Fig 4J-K and Sup Fig 4E). These data show that loss of En1 in the epithelium during regeneration does not affect UA digit bone length or volume but leads to a modest inhibition of bone regeneration.

We next bred Lmx1b mesenchymal-specific conditional knockout mice using an Lmx1b-flox allele and the Msx1-creERT2 tamoxifen inducible cre allele (Lopes et al., 2011; Zhao et al., 2006). To determine the role of Lmx1b in digit tip regeneration, we induced recombination in Lmx1b-fl/fl;Msx1-creERT2 mice with three daily doses of tamoxifen while leaving Lmx1b-fl/fl and Lmx1b-fl/fl;Msx1-creERT2 control cohorts uninduced. Tamoxifen dosing was sufficient to induce recombination of the Lmx1b locus and no ectopic recombination was detected in the control cohorts (Sup Fig 4F). We collected regenerated digit tips at 4wpa or 6wpa and performed uCT analyses to determine bone length and volume.

Gross qualitative assessment of the uCT scans showed that both the Lmx1b-fl/fl;Msx1-creERT2 cKO and Lmx1b-fl/fl control digit tip bones regenerated (Fig 4G, 4I). Quantitative analysis showed that the Lmx1b-fl/fl control regenerated digit tips are longer at 4wpa (p = 0.024) than contralateral UA controls, which is consistent with over-regeneration of the En1 control digits and previous literature (Fig 4J, 4L) (Dawson et al., 2018; Fernando et al., 2011). Uninduced Lmx1b-fl/fl;Msx1-creERT2 digits regenerate to a length equivalent to Lmx1b-fl/fl control digits, while Lmx1b-fl/fl;Msx1-creERT2 cKO regenerated digits are shorter at both 4wpa (p = 1.0×10^-7^) and 6wpa (p = 8.6×10^-4^) compared to Lmx1b-fl/fl control digits (Fig 4L). Lmx1b-fl/fl;Msx1-creERT2 cKO UA digits are also shorter than Lmx1b-fl/fl control UA digits (p = 0.049), implying that loss of Lmx1b may disrupt bone homeostasis. In terms of volume, Lmx1b-fl/fl control digits have a larger volume at 4wpa (p < 1×10-7) and 6wpa (p < 1×10-7) than UA, as expected and consistent with over-regeneration (Fig 4M). Uninduced Lmx1b-fl/fl;Msx1-creERT2 digits at 4wpa have not regenerated to the Lmx1b-fl/fl control volume (p = 0.010), but catch up by 6wpa, while Lmx1b-fl/fl;Msx1-creERT2 cKO digits have less volume at both 4wpa (p = 3.7×10^-5^) and 6wpa (p = 6.1×10^-5^) compared to Lmx1b-fl/fl controls (Fig 4M). These data indicate that mesenchymal Lmx1b is not required for digit tip bone regeneration, but loss of Lmx1b during regeneration does modestly impair bone regeneration. Therefore, the loss of regenerative ability in the Lmx1b developmental knockout (Lmx1b-fl/fl;Prx1-cre, Fig 3) can be attributed to the pre-existing digit dysmorphology, not the necessity of Lmx1b for regeneration.

### En1 and Lmx1b do not regulate dorsal-ventral bone patterning during digit tip regeneration

While our data find that Lmx1b and En1 are not genetically necessary for successful digit tip regeneration, we sought to determine if they are necessary for patterning the DV axis of the regenerating digit tip. The loss of En1 during limb development results in a “double dorsal” digit tip bone with a conical morphology (Fig 3E and 5A). To quantify the circularity of this conical morphology, we calculated the aspect ratio of uCT sections transecting the DV axis. The ratio of bone width and height were measured in the uCT sections, where an aspect ratio of one reflects a more circular shape (Fig 5A). It follows that uCT sections through the length of an En1-fl/fl;K14-cre neonatal digit tip bone have aspect ratios closer to 1 (Fig 5B-C) than comparable sections through an En1-fl/fl control digit tip bone (Fig 5D-E). The average aspect ratio of En1-fl/fl;K14-cre developmental knockout UA (0.86 ± 0.10) and regenerated (0.86 ± 0.10) digit tip bones are near one throughout the length of the digit tip, with no significant difference in ratio among the slices (Fig 5C). No significant difference in the aspect ratio was found between the En1-fl/fl;K14-cre UA and regenerated digit uCT slices, supporting our 2D alizarin red assessment (Fig 3F) that the regenerated digit maintains the double dorsal cylindrical morphology of the UA digit (Fig 5C).

**Figure 5.**
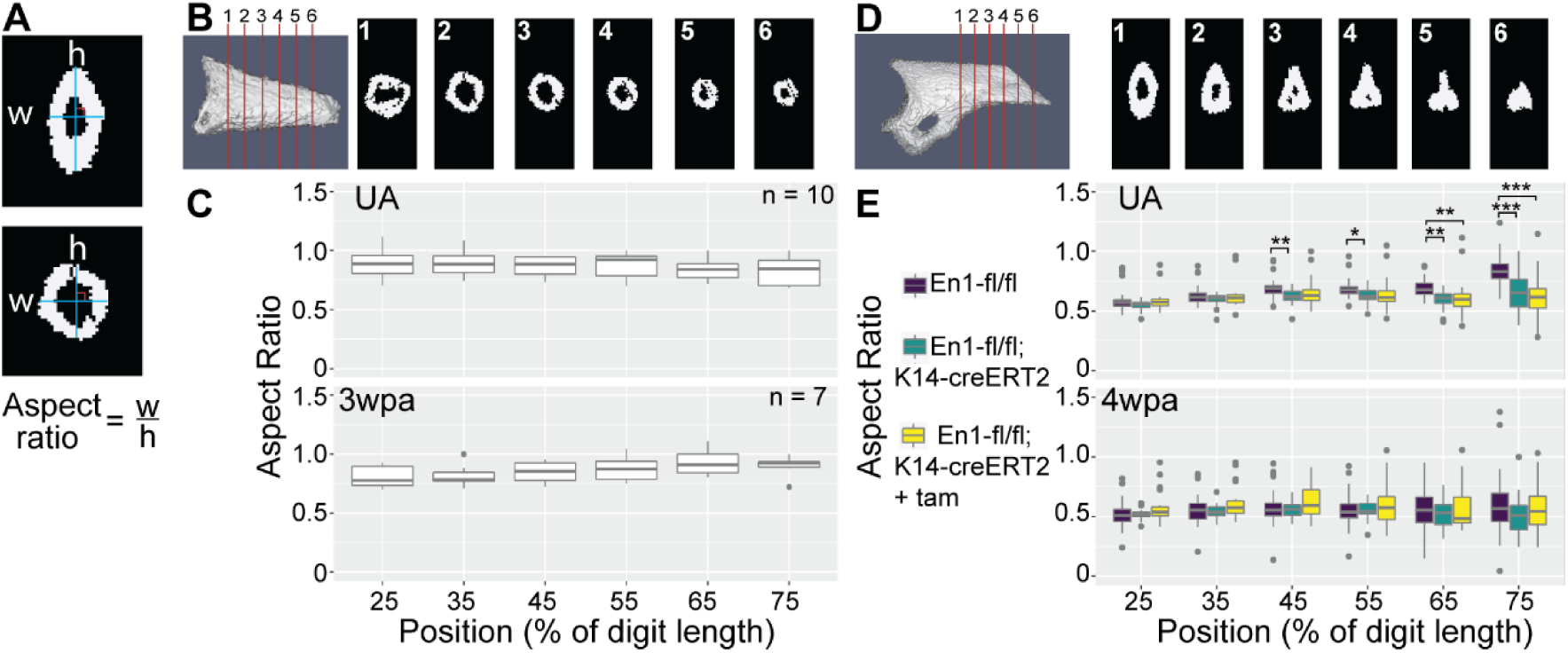
Loss of En1 during development, but not regeneration, increases circularity of the digit tip bone. (A) Example width and height measurements for particular slices shown in blue. (B) Example 3D rendering of a neonatal En1-fl/fl;K14-cre digit. Numbered red lines indicate approximate position of correspondingly numbered slices through the digit bone. (C) Box plot of aspect ratio measure for slices at 25, 35, 45, 55, 65, and 75 percent of the length of the digit in unamputated (top) and 3wpa (bottom) digits. n refers to the number of individual digits in each group. (D) Example 3D rendering of an adult En1-fl/fl digit. Numbered red lines indicate approximate position of correspondingly numbered slices through the digit bone. (E) Box plot of aspect ratio measure for slices at 25, 35, 45, 55, 65, and 75 percent of the length of the digit starting from the distal most part of the ventral hole in unamputated (top) and 4wpa (bottom) digits. UA En1-fl/fl n = 48, UA En1-fl/fl;K14-creERT2 n = 24, UA En1-fl/fl;K14-creERT2 + tam n = 28. 4wpa En1-fl/fl n = 48, 4wpa En1-fl/fl;K14-creERT2 n = 22, 4wpa En1-fl/fl;K14-creERT2 + tam n = 27. n refers to the number of individual digits in each group. * p < 0.05, ** p < 0.01, *** p < 0.001.

In the adult En1 cKO model, UA En1-fl/fl control (0.68 ± 0.11), uninduced En1-fl/fl;K14-creERT2 (0.61 ± 0.09), and En1-fl/fl;K14-creERT2 cKO (0.61 ± 0.13) digits have smaller aspect ratios than the En1-fl/fl;K14-cre developmental knockout, consistent with non-circular morphology. Despite variability in the aspect ratio measurement at the tip of the UA digit, En1-fl/fl digits have larger aspect ratios than En1-fl/fl;K14-creERT2 digits at 45 (p = 0.006), 55 (p = 0.042), 65 (p = 0.002), and 75% (p = 8.1×10^-6^) of the digit tip length, and En1-fl/fl;K14-creERT2 plus tamoxifen digits at 65 (p = 0.002) and 75% (p < 1×10-7) of digit length (Fig 5E). This likely reflects stochasticity in the shape of the digit bone near the tip. If En1 regulates DV patterning during regeneration, we would expect the regenerated bone in En1-fl/fl;K14-creERT2 cKO digits to have a circular, double dorsal, morphology. The average aspect ratios for these digit uCT slices are similar to regenerated En1-fl/fl control (0.56 ± 0.15), uninduced En1-fl/fl;K14-creERT2 (0.53 ± 0.10), and En1-fl/fl;K14-cre-ERT2 cKO (0.60 ± 0.16) digits, with no significant differences in aspect ratio at any slice (Fig 5E bottom). Overall, this analysis shows that deletion of En1 during regeneration does not lead to a circular, double dorsal bone morphology, and thus En1 does not specify ventral fate during digit tip regeneration.

More generally, loss of En1 and Lmx1b during limb development result in near perfect DV symmetry throughout the digit tip bone (Fig 3E and 3K). If En1 or Lmx1b are involved in DV patterning of the regenerating digit tip, then loss of either during regeneration should increase DV symmetry in the regenerated digit bone. To measure DV symmetry of the digit tip bone, we placed landmarks at the most dorsal and most ventral points on a fixed slice on each digit tip bone uCT scan (Fig 6A, red cubes) and used this index slice to bisect the digit into dorsal and ventral compartments (Fig 6A). We measured the dorsal and ventral areas progressively from the index slice to the distal tip, computed the ventral to dorsal area ratio as a measure of symmetry (Fig 6A), and plotted the ratio for each slice as a function of normalized position along the digit tip bone (Sup Fig 5A, 5B). The resulting curve shows a stereotyped shape where segment 1 (s1, Fig 6A) corresponds to the morphology of the proximal digit, the change point corresponds to where the ventral volume of the digit stops decreasing (which is also where the amputation plane is located) (cp, Fig 6A), and the second segment corresponds to the morphology of the regenerated part of the digit when amputated (s2, Fig 6A). To determine if these parameters significantly change after regeneration in the En1 or Lmx1b cKO models, we used the mcp R package (Lindeløv, 2020) to fit a segmented linear regression model for each digit. The resulting model estimates parameters that describe the slope of s1, the change point position, and the slope of s2.

**Figure 6.**
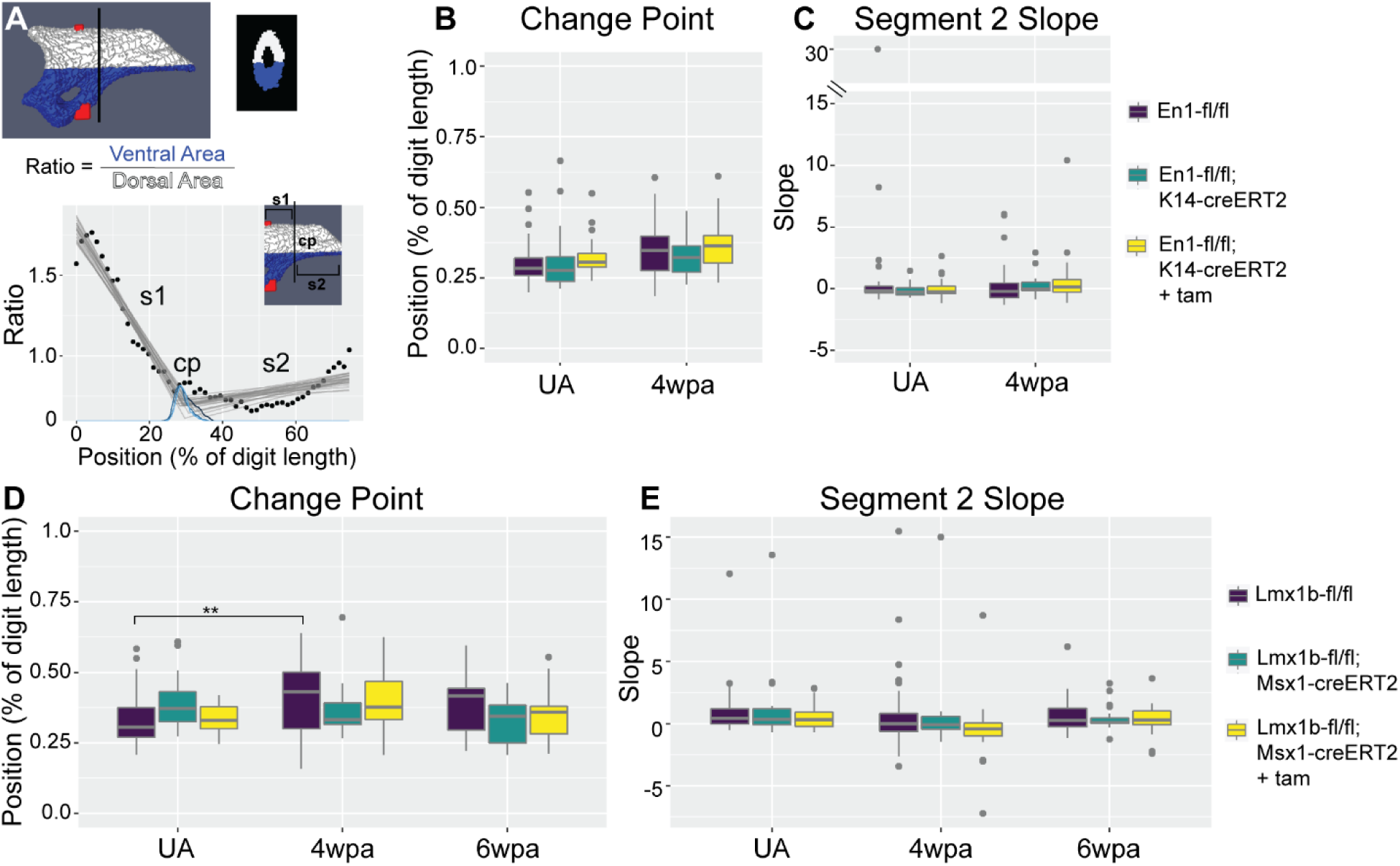
Conditional knockout of En1 or Lmx1b does not affect overall morphology of regenerated digit tip bones. (A) Schematic of the concept behind data and analysis in (B-E). An unamputated digit is shown with dorsal area in white and ventral area in blue (top left), with a slice at the approximate position of the black line (top right). A ratio is computed by taking the ventral area divided by the dorsal area at many slices along the length of the digit. The plotted ratios (black points) are within the area shown by the inset, with the fitted model shown as grey lines. s1 = first segment, cp = change point, s2 = second segment. (B) Box plot showing mean change point predictions for each digit in the En1 cohort. (C) Box plot showing mean slope predictions for the second segment for each digit in the En1 cohort. (D) Box plot showing mean change point predictions for each digit in the Lmx1b cohort. (E) Box plot showing mean slope predictions for the second spline segment in the Lmx1b cohort. Number of individual digit tip bones in each group is noted in methods and sup fig 5. * p < 0.05, ** p < 0.01, *** p < 0.001.

After fitting a segmented line to each individual digit, we compared the biologically meaningful parameters between groups to determine if the DV morphology of the bone is changed. We find that En1-fl/fl;K14-creERT2 cKO digit tips are not significantly different than En1-fl/fl control UA digits in either change point position or the slope of segment 2 (MANOVA p = 0.48) (Fig 6B, C), illustrating that loss of En1 in the epithelium does not change the morphology of the digit tip bone during homeostasis. Furthermore, the change points and segment 2 slopes of 4wpa regenerated En1-fl/fl;K14-creERT2 cKO digits are not significantly different from En1-fl/fl control 4wpa digits; the same is true for uninduced, but leaky, En1-fl/fl;K14-creERT2 digits at both UA and 4wpa time points (MANOVA p = 0.48) (Fig 6B-C). This suggests that loss of En1 in the epithelium during regeneration does not change the morphology or DV symmetry of the regenerated digit tip, further underscoring that En1 does not participate in DV patterning during digit tip regeneration.

We performed the same DV symmetry analysis for the Lmx1b cKO mutants and controls. No significant difference was found in change point position or segment 2 slope between Lmx1b-fl/fl control and Lmx1b-fl/fl;Msx1-creERT2 uninduced UA digits (cp p = 0.21, s2 slope ANOVA p = 0.59). The same was true when comparing Lmx1b-fl/fl control and Lmx1b-fl/fl;Msx1-creERT2 cKO (cp p = 1, s2 slope ANOVA p = 0.59) UA digits, which indicates that loss of Lmx1b does not change homeostatic digit morphology (Fig 6D-E). Similarly, at 4wpa, Lmx1b-fl/fl;Msx1-creERT2 uninduced and cKO digits have no difference in either change point (p = 0.97 and p = 0.99, respectively) or segment 2 slope (ANOVA p = 0.59) compared to Lmx1b-fl/fl 4wpa controls (Fig 6D-E). By 6wpa, Lmx1b-fl/fl;Msx1-creERT2 uninduced and cKO digits also do not have significantly different change point (p = 0.61 and p = 0.99, respectively) or segment 2 slope (ANOVA p = 0.59) values compared to Lmx1b-fl/fl controls (Fig 6D-E), indicating that loss of Lmx1b does not change DV morphology in the UA or regenerated digit tip. Interestingly, the segment 1 slope, which corresponds to morphology of the proximal digit, is significantly increased at 4wpa (p < 1×10-7) and 6wpa (p = 0.007) compared to UA for both Lmx1b-fl/fl;Msx1-creERT2 cKO digits (p=0.00 and p=1.2×10^-4^, respectively) as well as Lmx1b-fl/fl control digits (p=0.00 and p=0.007, respectively) (Sup Fig 5C). This suggests both the control and cKO regenerated digits have more ventral bone proximal to the amputation plane in the regenerated digits, but this phenotype is not Lmx1b specific. We hypothesize that the mixed background strains of the Lmx1b-flox and Msx1-creERT2 alleles may promote extended histolysis following amputation, which would cause bone regeneration to also occur more proximally (Lehoczky et al., 2011; Sammarco et al., 2015). The increase in the measured change points of Lmx1b-fl/fl digits at 4wpa compared to UA (p = 0.004) could be caused by an increase in uncertainty in modeling the change point because the slopes of s1 and s2 are more similar (Sup Fig 5B). Overall, we find that loss of Lmx1b in the mesenchyme during regeneration does not affect the DV symmetry of the digit tip.

## DISCUSSION

Recapitulation of developmental gene expression has long been hypothesized to be the mechanism of pattern formation during regeneration (Goss, 1969). We tested this hypothesis by investigating whether En1 and Lmx1b, two transcription factors necessary for DV patterning during limb development, are also necessary for DV patterning during mouse digit tip regeneration. We show that En1 and Lmx1b are expressed during digit tip regeneration but not with DV polarity. Furthermore, with conditional deletion during regeneration, we show that En1 and Lmx1b are not genetically necessary for DV patterning during mouse digit tip regeneration. This is unexpected given that in other models of regeneration, particularly zebrafish fin or salamander limb regeneration, developmental patterning genes are expressed in domains similar to development (Gardiner et al., 1995; Han et al., 2001; Rabinowitz et al., 2017; Roensch et al., 2013; Shimokawa et al., 2013; Torok et al., 1999).

If not recapitulating the developmental patterning function, what is the function of En1 or Lmx1b during digit homeostasis and regeneration? Our finding that deletion of En1 in the epithelium during homeostasis results in the ventral toe pad transforming to a dorsal nail-like structure (En1-fl/fl;K14-creERT2; Sup Fig 4B), suggests that En1 continues to play a role in maintaining ventral identity in the epithelium and ectodermal appendages of the adult homeostatic digit. However, since En1 is expressed in both ventral and dorsal epithelium in the adult mouse, which is distinct from limb development, another factor in the epithelium may modulate the maintenance of ventral identity by En1. Beyond DV identity, our data suggest that En1 expression in the epithelium supports bone regeneration because En1-fl/fl;K14-creERT2 regenerated digits had a modest reduction in bone length and volume.

Digit tip regeneration is closely linked to the nail: amputation proximal to the nail fails to regenerate, and Wnt signaling from the nail bed promotes regeneration (Lehoczky and Tabin, 2015; Mohammad et al., 1999; Takeo et al., 2013; Zhao and Neufeld, 1995). Here we show that En1-fl/fl;K14-cre regenerated digit tips, with nails that encircle the entire digit, are longer than regenerated controls (Fig 3M). This is not seen in En1-fl/fl;K14-creERT2 mice, which have a slight reduction of bone length after regeneration (Fig 4J). These seemingly opposite regenerative responses may be attributed to differences in the nail, which is circumferential in En1-fl/fl;K14-cre mice and normal in En1-fl/fl;K14-creERT2 mice. It follows that the nail epithelium may provide a pro-regenerative signals or the nail plate itself may be integral to patterning the bone during digit tip regeneration by exerting a mechanical force on the tissue (Johnson and Lehoczky, 2021; Lehoczky, 2017).

The role of Lmx1b in digit homeostasis and regeneration is less clear. During homeostasis Lmx1b expression is restricted to the dorsal mesenchyme, which lies beneath the nail epithelium. The nail has been shown to modulate digit tip regeneration partly through Wnt signaling (Lehoczky and Tabin, 2015; Takeo et al., 2013). During limb development, Lmx1b is downstream of Wnt7a. While Wnt7a is not expressed in the epithelium of homeostatic or regenerating digit tips (Sup Fig 3), another Wnt ligand expressed in the epithelium may induce Lmx1b expression in the digit tip mesenchyme. The developmental function of Lmx1b suggests that it could function in maintaining the dorsal identity of the nail epithelium during homeostasis, though we do not find transformation or loss of the nail in our Lmx1b conditional mutants up to 4 weeks after tamoxifen administration.

Furthermore, we find that conditional deletion of Lmx1b results in a modest reduction in bone regeneration. In contrast, Lmx1b has previously been characterized as anti-osteogenic during calvarial development, where Lmx1b is expressed in a population of mesenchymal cells that reside in the suture and loss of Lmx1b results in premature suture closure (Cesario et al., 2018). Lmx1b is also expressed in the joint mesenchyme surrounding the bone during mouse limb development and zebrafish fin regeneration (Dreyer et al., 2004; Tang et al., 2022). While our results seem contrary to these findings, it is possible that Lmx1b has a similar anti-osteogenic function in mouse digit tip regeneration. Loss of Lmx1b in blastema cells could cause premature osteogenic differentiation during regeneration, thereby disrupting regeneration and causing loss of regenerated bone length and volume. Alternatively, Lmx1b could have a separate function in the context of adult digit tip homeostasis and regeneration. Loss of Lmx1b also results in shorter digit tip bones independent of regeneration, pointing to a general role in bone homeostasis and maintenance.

By visual inspection, the imperfect regeneration of the digit tip bone could be interpreted as evidence that there is no pre-determined pattern that forms during regeneration. However, our uCT analysis shows that there is a reproducible DV pattern in the regenerated digit tip bone. Here we present data that show developmental DV patterning genetic networks are not re-used during adult digit tip regeneration, which suggests that other factors regulate DV patterning during regeneration. It is difficult to determine candidates for other factors based on limb development alone because effectors of DV pattern downstream of En1 are largely unknown. ChIP-seq for targets of Lmx1b during limb development has identified many regulatory targets of Lmx1b, but how these genes drive morphological differences is not understood (Haro et al., 2017). That said, expression of Lmx1b throughout the blastema could suggest that the blastema is an entirely dorsal structure. However, other experimental evidence does not support this idea. Hyperbaric oxygen treatment causes degradation of the digit tip bone almost entirely after amputation in some cases (Sammarco et al., 2015). If the blastema were entirely dorsal, the regenerated bone would not form ventral structures like the ventral hole. However, the regenerated bone correctly re-patterns the ventral hole, implying that dorsal and ventral identities are conserved in the blastema during regeneration (Sammarco et al., 2015).

Collectively our data demonstrate that at least one developmental limb patterning network is not re-used to regulate patterning during digit tip regeneration. However, En1 and Lmx1b only define the DV axis during limb development and there are two other axes (AP and PD) to consider. The digit tip bone is symmetrical in the AP axis, while there are morphological changes along the PD axis. Dermal fibroblasts, an important population in digit tip regeneration, retain positional information in the form of location specific Hox expression, providing a Hox dependent mechanism of positional memory (Rinn et al., 2006; Rinn et al., 2008). Therefore, Hoxa13 and Hoxd13 are promising candidates that may direct morphogenesis in the PD and/or AP axes during digit tip regeneration as they do during limb development (Dollé et al., 1993; Fromental-Ramain et al., 1996; Kmita et al., 2005). In contrast to loss of En1 or Lmx1b during development, which results in a quantifiable digit tip bone phenotype, loss of Hoxa13 or Hoxd13 during development do not result in a morphological change in the digit tip P3 bone but instead loss of certain digits, loss of the P2 bone, and delayed development of digits overall (Fromental-Ramain et al., 1996). However, Hoxd13 and Hoxa13 can also induce expression of osteogenic genes like Runx2, Bmp2, and Bmp7, coupling patterning and bone formation (Knosp et al., 2004; Takeo et al., 2013; Yu et al., 2010) and complicating interpretation of their role in regenerate patterning. By focusing on the DV axis in our study, we are able to distinguish between changes in bone regeneration and patterning, and between expression and function of developmental patterning genes during regeneration. Collectively, our data support the involvement of non-developmental mechanisms in patterning the mouse digit tip during regeneration. It is important to determine these patterning mechanisms because inducing regeneration in non-regenerative contexts, such as human limb regeneration, will require correct patterning of the regenerate.

## MATERIALS AND METHODS

### Mouse husbandry and surgery

Mouse (*Mus musculus*) husbandry and surgeries were done with the approval of the Brigham and Women’s Hospital IACUC. CD1(ICR) mice used for e10.5 hindlimb, e12.5 brain, and fetal and neonatal digit tip tissue were obtained from Charles River Laboratories (Cat #CRL:022). FVB/NJ mice used for adult regenerating and unamputated digit tip tissue were obtained from The Jackson Laboratory (Cat #JAX:001800). To achieve En1 tissue specific knockout during development an En1 floxed allele (Sgaier et al., 2007) (Cat #JAX:007918) was used in combination with a K14-Cre allele (Dassule et al., 2000) (Cat #JAX:004782). En1 conditional knockout studies in adults used the same En1 floxed allele in combination with a K14-creERT2 allele (Li et al., 2000) (courtesy of Dr. Pierre Chambon). To achieve Lmx1b mesenchymal specific knockout during development an Lmx1b floxed allele (Zhao et al., 2006) (courtesy of Dr. Randy Johnson) was used in combination with a Prx1-Cre allele (Logan et al., 2002) (Cat #JAX:005584). Lmx1b conditional knockout studies in adults used the same Lmx1b floxed allele in combination with an Msx1-CreERT2 allele (Lopes et al., 2011) (courtesy of Dr. Benoit Robert). To induce recombination in 4-week-old En1-fl;K14-CreERT2 and 6-week-old Lmx1b-fl;Msx1-CreERT2 mice, tamoxifen (Sigma-Aldrich #T5648-1G) in corn oil was administered by oral gavage at 3mg/40g mouse body weight. Daily doses were administered on three consecutive days and digits were amputated the day after the last dose.

Neonatal digit amputations were performed for En1-flox;K14-Cre and Lmx1b-flox;Prx1-Cre experimental cohorts as previously described (Lehoczky and Tabin, 2015; Lehoczky et al., 2011). Postnatal day 3 pups were cryoanesthetized and hindlimb digits 2, 3, and 4 were amputated midway through the distal tip with micro-spring scissors. Pups were returned to the mother and digits regenerated until harvested as noted. Adult digit tip amputations were performed as previously reported (Johnson et al., 2020). FVB/NJ, En1-flox;K14-CreERT2, or Lmx1b fl; MsxCreERT2 mice were anesthetized with 3-4% isoflurane and anesthesia was maintained at 2% isoflurane. Digits 2, 3, and 4 of the right and/or left hindlimb were visualized under a Leica S6E dissecting microscope and amputated distally. Mice were given buprenorphine at a dose of 0.05 mg/kg at time of surgery and again 8-12 hours later, then monitored for 4 days.

### Hybridization chain reaction RNA fluorescent in situ hybridization (HCR RNA-FISH)

Unamputated digit tissue from neonatal (PN4/5) CD1(ICR) mice was collected into ice cold 4% PFA for fixation overnight at 4C, washed with PBS, and then incubated in graded sucrose solutions from 5% to 30% over 3 days at 4C before embedding in OCT (Tissue-Tek). Unamputated embryonic (e10.5 or e12.5) and fetal (e16.5) digit tissue was collected from timed pregnant CD1(ICR) females and treated as above with solution changes at 30 minutes instead of overnight. Adult unamputated or blastema stage digits from FVB/NJ, En1-flox;K14-creERT2, or Lmx1b-flox;Msx1-creERT2 mice were collected and treated in the same manner as neonatal digit tissue with the addition of a decalcifying step with Decalcifying Solution Lite (Sigma-Aldrich) at room temperature for 40 minutes before graded sucrose. All tissue was stored in OCT at -80C and then sectioned at 10μm (fetal and neonatal) or 18μm (adult) on a Leica CM3050S cryostat.

For HCR RNA-FISH, sections were prepared for probe hybridization as previously described (Murtaugh et al., 1999) with the addition of Proteinase K (3ug/mL) for 10 minutes at room temperature. Probe hybridization and signal amplification followed the Molecular Instruments HCR v3.0 manufacturer’s protocol for sections. Briefly, probes targeting mouse En1 (NM_010133.2), Lmx1b (NM_010725.3), and Wnt7a (NM_009527.4) were hybridized to tissue sections at 37C overnight. After probe washing, signal amplification hairpins were added to sections and amplified at room temperature overnight. After washing, the TrueVIEW autofluorescence quenching kit (Vector Laboratories) was used to quench endogenous fluorescence before sections were counterstained with DAPI at 1ng/uL. To generate no probe control sections, sections were treated as above but no probe was added to the hybridization buffer. All in situ experiments were done minimally in duplicate on sections from different animals. Sections were imaged in 5-10uM z-stacks on a Zeiss LSM 880 confocal microscope at 20x or 40x. Z-stack maximum projections were generated in ImageJ v.1.53c and brightness and contrast of pseudo colored magenta, green, and gray channels were adjusted independently in Adobe Photoshop v.22.0.0.

### Skeletal staining and analysis

3wpa and control UA digits of Lmx1b-flox;Prx1-cre and En1-flox;K14-cre experimental cohorts were prepared with a standard alizarin red and alcian blue staining protocol as previously reported (Lehoczky and Tabin, 2015; McLeod, 1980). Briefly, digits were collected into 100% ethanol for dehydration, then incubated in staining solution (0.005% alizarin red, 0.015% alcian blue, 5% acetic acid, and 60% ethanol) at 37°C for 24 hours. Digit tissue was cleared using a 2% potassium hydroxide at room temperature for 2 days, then taken through increasing concentrations of glycerol (25%, 50%, 75%, and 100%) and imaged on a Leica M165 FC stereo microscope with a Leica DFC7000 T camera attachment. Images were processed in ImageJ where length was manually measured in triplicate from the mid-proximal joint to the distal digit tip. Regenerated digit length was divided by the same animal’s contralateral UA digit length to calculate percent regeneration. In total, 15 En1-wt/wt;K14-cre, 8 En1-fl/fl;K14-cre, 15 Lmx1b-fl/wt;Prx1-cre and 6 Lmx1b-fl/fl;Prx1-cre digits were analyzed. Data are represented using boxplots plotted in R (v4.1.2) (R Core Team, 2018) using the ggplot2 package (Wickham, 2016) where the center line corresponds to the median value, upper and lower edges of the box denote the 75^th^ and 25^th^ percentile values respectively, and the whiskers extend to the smallest or largest value within 1.5 times of the interquartile range. A two-tailed t-test was used to determine statistical significance between groups.

### Micro computed tomography (uCT) scanning and analysis

4wpa or UA digits from Lmx1b-flox;Msx1-creERT2 and En1-flox;K14-creERT2 experimental cohorts were collected and stored in 70% ethanol until scanning. A Scanco Medical uCT 35 system with an isotropic voxel size of 7um was used to scan digits in 70% ethanol in a 7mm diameter sample tube with an X-ray tube potential of 55 kVp, a 0.5 mm aluminum filter, an X-ray intensity of 0.145 mA, and an integration time of 600 ms per slice. Alizarin red stained 3wpa or UA developmental mutant digits from En1-flox;K14-cre mice were scanned in 25% glycerol with the same settings. Scans were then converted to DICOM image sets and imported into our custom image analysis software, which used the C programming language for low level functions and the IDL package (L3Harris Geospatial, Broomfield, CO) for the graphical user interface. This semi-automated image analysis software was based on a variable thresholding algorithm and modeled on previous software tools to measure carpal bone volume for rheumatoid arthritis assessment (Duryea et al., 2008) and to quantify zebrafish vertebral morphometry (Charles et al., 2017). We first placed two seed points on the digit scans identifying the base and tip of the distal phalanx. This step was used both to identify the location of the distal phalanx on the scan and to determine a “bone axis”, defined as the approximate direction of the digit. Next, we chose a grey scale threshold level optimized to isolate the distal phalanx from the other more proximal bones in the scan, and the total bone volume and length were calculated. The isolated digit tip bone was then rotated so that the bone axis was aligned with one edge of the 3D image so we could view cross sections of the distal phalanx. Landmarks were then placed on the rotated 3D images, one marking the slice containing the most distal tip of the bone and the other at an indexed slice located at the most distal end of the ventral hole. Two landmarks were placed on the indexed slice to define the DV axis (Fig 6A, red points) from which two additional morphometric metrics were defined. A symmetry value was defined as the ratio of the areas above and below the midpoint axis, which was placed at 60% of the height of the digit at the index slice. This ratio was plotted for the length of the bone from the indexed slice to the end point (Fig 6). We also defined a measure of the height to width, referred to as an aspect ratio, which was calculated at fixed locations along the digit (Fig 5A). The previously defined DV axis at the index slice was used to measure the height of the bone at each slice, then a line perpendicular to the first line at 50% of the height was placed and the bone width measured. The aspect ratio was calculated as width divided by height. To fit segmented line (spline) models to each digit, we utilized the mcp package (v0.3.1) in R to infer two segments and one change point for each digit (Lindeløv, 2020). Ratio values for 0 to 75% of digit length were used as input to the model. We disregarded ratio values for the last 25% of the digit because morphology at the tip is variable, leading to ratio values of 0 or infinity for many digits.

Sample sizes for each group used in the length, volume, and spline analyses are as follows, where n refers to the number of individual digits in each group. UA: En1-fl/fl n = 48, En1-fl/fl;K14-creERT2 n = 24, En1-fl/fl;K14-creERT2 + tam n = 28, Lmx1b-fl/fl n = 60, Lmx1b-fl/fl;Msx1-creERT2 n = 34, Lmx1b-fl/fl;Msx1-creERT2 + tam n = 45. 4wpa: En1-fl/fl n = 48, En1-fl/fl;K14-creERT2 n = 22, En1-fl/fl;K14-creERT2 + tam n = 27, Lmx1b-fl/fl n = 39, Lmx1b-fl/fl;Msx1-creERT2 n = 13, Lmx1b-fl/fl;Msx1-creERT2 + tam n = 27. 6wpa: Lmx1b-fl/fl n = 23, Lmx1b-fl/fl;Msx1-creERT2 n = 23, Lmx1b-fl/fl;Msx1-creERT2 + tam n = 14. For the circularity analysis, En1-fl/fl;K14-cre UA n = 10, En1-fl/fl;K14-cre 3wpa n = 7, UA En1-fl/fl n = 48, UA En1-fl/fl;K14-creERT2 n = 24, UA En1-fl/fl;K14-creERT2 + tam n = 28, 4wpa En1-fl/fl n = 48, 4wpa En1-fl/fl;K14-creERT2 n = 22, 4wpa En1-fl/fl;K14-creERT2 + tam n = 27.

To test for statistically significant differences between groups for digit length, volume, and section circularity in R (v4.1.2) (R Core Team, 2018), significant two-way ANOVA tests were followed by a post-hoc Tukey HSD test with multi comparison correction. Significant two-way MANOVA tests were followed with a two-way ANOVA and, if warranted, a Tukey HSD post-hoc test with multiple comparison correction to test for statistically significant differences between groups for mean spline parameters. Adjusted p-values are reported. Significant p-values are denoted as * if p < 0.05, ** if p < 0.01, and *** if p < 0.001. Data are represented by boxplots as above, where the center line corresponds to the median value and individual dots represent outliers.

### Quantitative PCR (qPCR)

Digits 1 and 5 from experimental En1-flox;K14-CreERT2 and Lmx1b-flox;Msx1-CreERT2 animals were collected and stored at -20C until DNA extraction with the DNEasy Blood and Tissue kit (Qiagen) according to kit instructions. A standardized DNA concentration was used as input to qPCR reactions with primers that detect the En1 recombined allele (En1-fl;K14-CreERT2 cohort) or the Lmx1b recombined allele (Lmx1b-fl;Msx1-CreERT2 cohort). These primers span introns or intron-exon junctions and thus do not amplify En1 or Lmx1b mRNA. Primers for detecting recombination of the En1-floxed allele were 5’-GAGCTTGCGGAACCCTTAAT-3’ and 5’-GGTAGAGAAGAGGCGAGG-3’. Primers for detecting recombination of the Lmx1b-floxed allele were 5’-CAGCCCAATTCCGATCATATTCA-3’ and 5’-AAATACGGGGCTTTGGAACA-3’. SsoAdvanced Universal SYBR Green Supermix (Bio-Rad) was used to detect amplification of the target sequences on a QuantStudio 5 Real-Time PCR system (Applied Biosystems). Data are represented by boxplots as above, where the center line corresponds to the median value and individual dots represent outliers.

## ACKNOWLEDGEMENTS

We thank the NeuroTechnology Studio at Brigham and Women’s Hospital for providing Zeiss LSM 880 confocal access and consultation on data acquisition. We also thank Dr. Anne Golding for Masson’s trichrome histology of the En1 mutant digit. We are grateful to other members of the Lehoczky lab and Drs. Cliff Tabin, Marian Ros, and Kerby Oberg for helpful scientific discussions.

## AUTHOR CONTRIBUTIONS

GLJ, JFC, JD, and JAL conceptualized and designed experiments. GLJ, MBG, and JAL performed experiments and acquired data. GLJ, MBG, JD, and JAL analyzed data. GLJ and JAL wrote the manuscript which was edited by all authors.

---

**Supplementary Figure 1.**
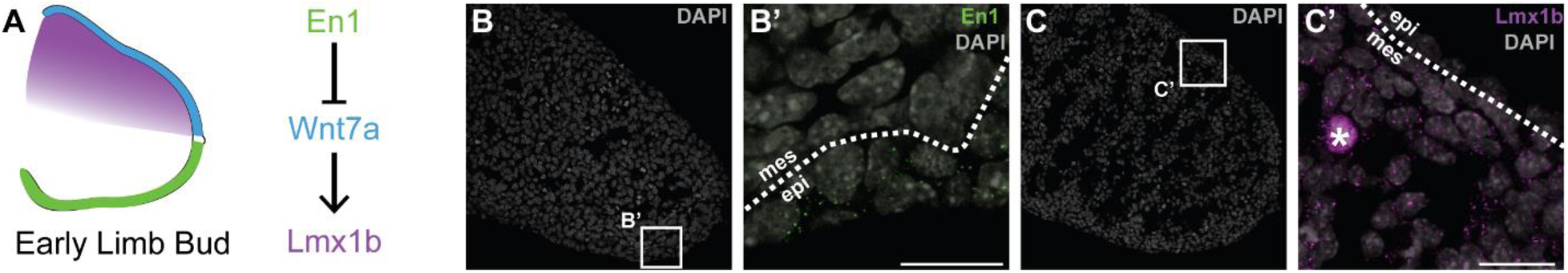
En1 and Lmx1b expression in the early limb bud. (A) Schematic of expression and interaction of En1, Lmx1b, and Wnt7a in the early limb bud. (B-C) HCR RNA-FISH in an e10.5 limb bud for (B-B’) En1 (green puncta) and (C-C’) Lmx1b (magenta puncta) with DAPI in grey. White boxes on DAPI only panels (B, C) show the position of magnified panels to the right. White dashed line outlines the epithelium. Scale bars represent 20μm. Asterisks (*) denote blood vessel autofluorescence. Epi = epithelium, mes = mesenchyme.

**Supplementary Figure 2.**
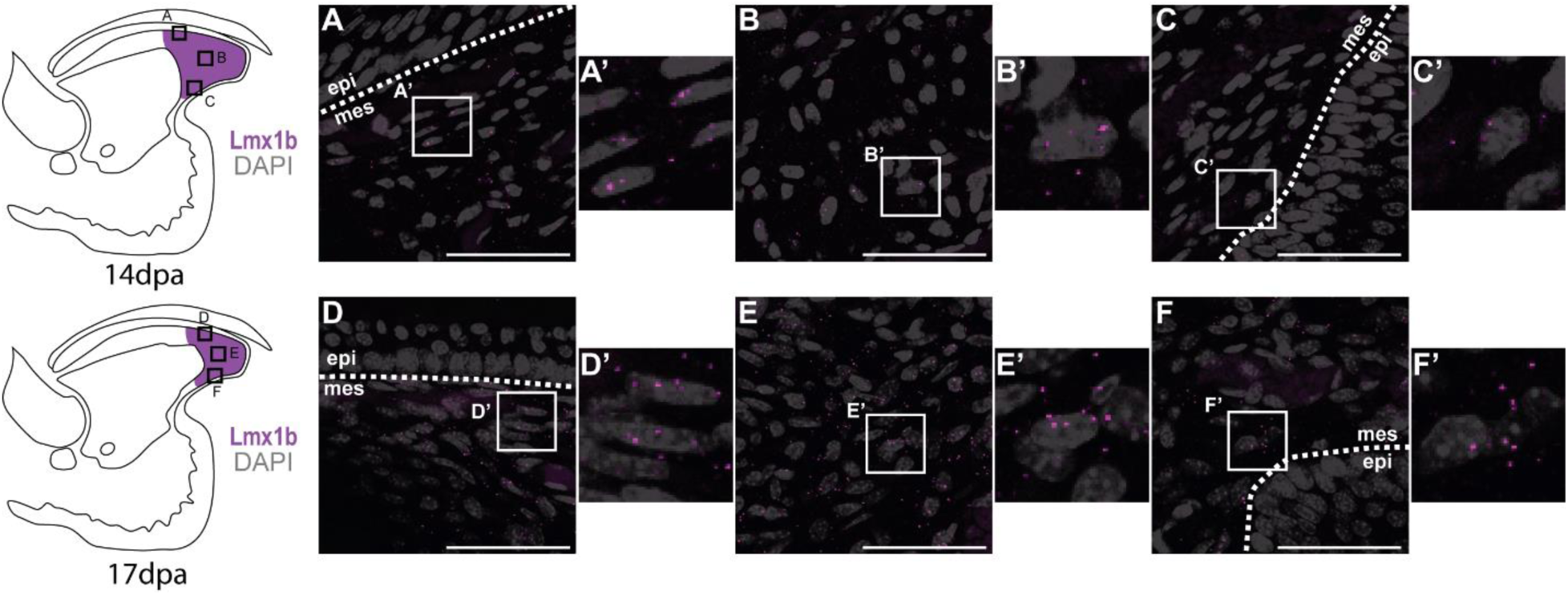
Lmx1b expression in later stages of regeneration. (A-F) HCR RNA-FISH for Lmx1b (magenta puncta) in (A-C) a 14dpa and (D-F) a 17dpa regenerating digit tip at dorsal, mid-blastemal, and ventral positions. Cartoons to the left show the approximate location of the corresponding lettered panel and include blastemal Lmx1b expression in magenta. White boxes show the position of magnified panels to the right. White dashed line outlines the epithelium. Scale bars represent 50μm. Epi = epithelium, mes = mesenchyme.

**Supplementary Figure 3.**
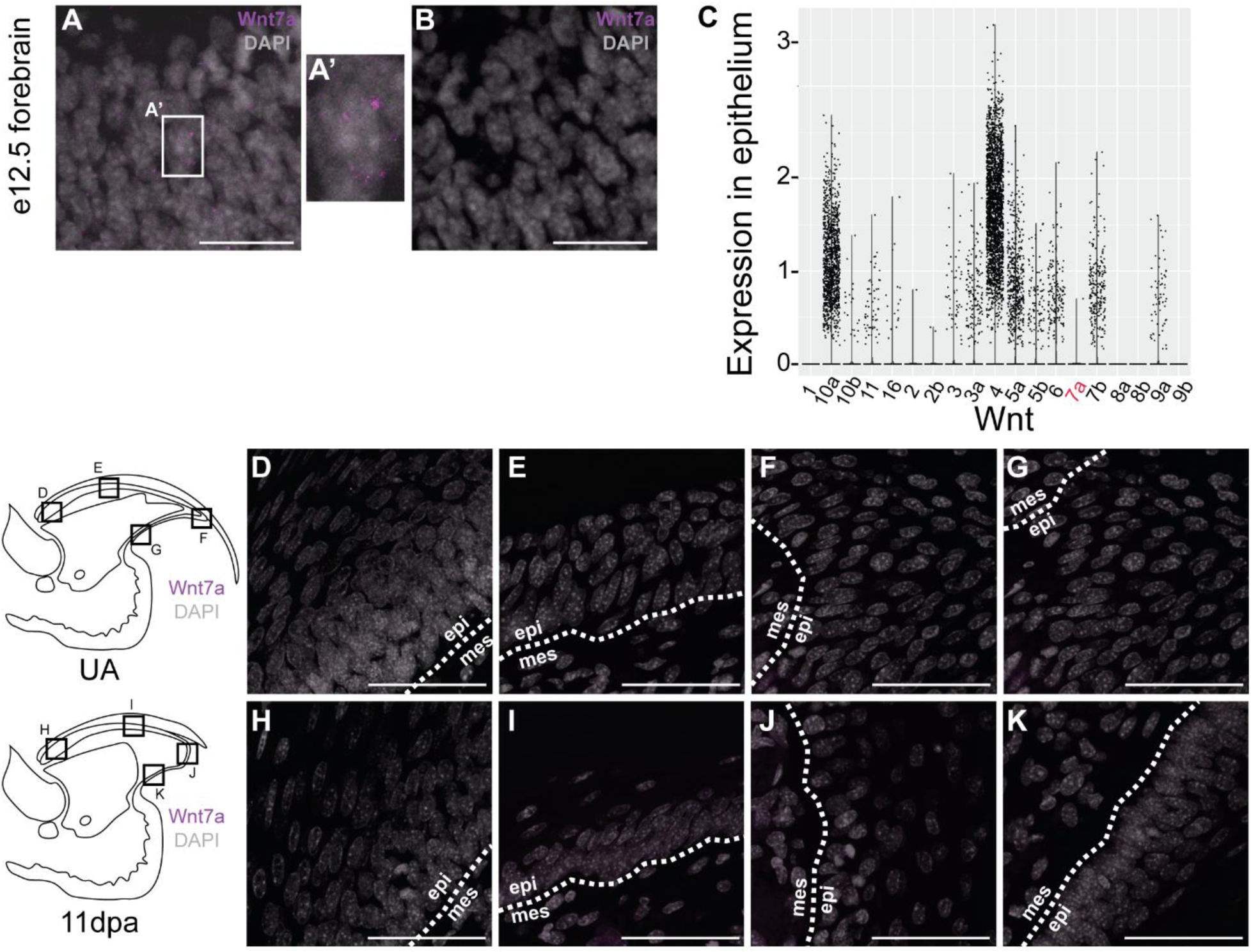
Wnt7a expression during digit tip homeostasis and regeneration. (A-B) HCR RNA-FISH for Wnt7a (magenta puncta) in mouse e12.5 forebrain sections. White box in (A) shows the position of the magnified panel (A’) to the right. (B) Shows a no probe control section. Scale bars represent 20μm and DAPI is shown in grey. (C) Violin plot of the expression of Wnt ligands in the epithelium in a combined set of UA, 11, 12, 14, and 17dpa cells. Dots represent individual cells. Wnt7a is highlighted in red. (D-K) HCR RNA-FISH for Wnt7a (magenta puncta) in a (D-G) UA and (H-K) 11dpa regenerating digit tip. Cartoons to the left show the approximate location of the corresponding lettered panel. White dashed lines outline the epithelium. Scale bars represent 50μm. Epi = epithelium, mes = mesenchyme.

**Supplementary Figure 4.**
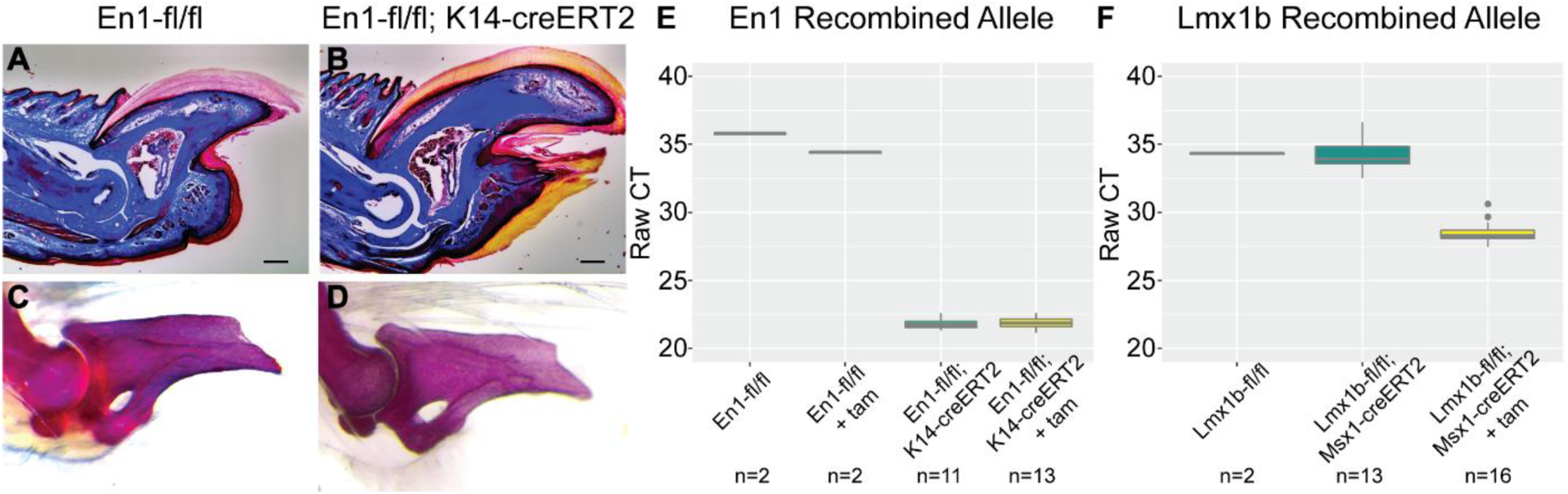
K14-creERT2 ectopic recombination of En1-floxed allele. (A-B) Masson’s trichrome stain of (A) En1-fl/fl control and (B) En1-fl/fl; K14-creERT2 UA digits. Scale bar represents 200μm. (C-D) Alizarin red skeletal stain of (C) En1-fl/fl control and (D) En1-fl/fl; K14-creERT2 UA digits. (E-F) Box plot of raw CT values from qPCR targeting the (E) En1-fl recombined allele or (F) Lmx1b-fl recombined allele. n refers to the number of individual digits.

**Supplementary Figure 5.**
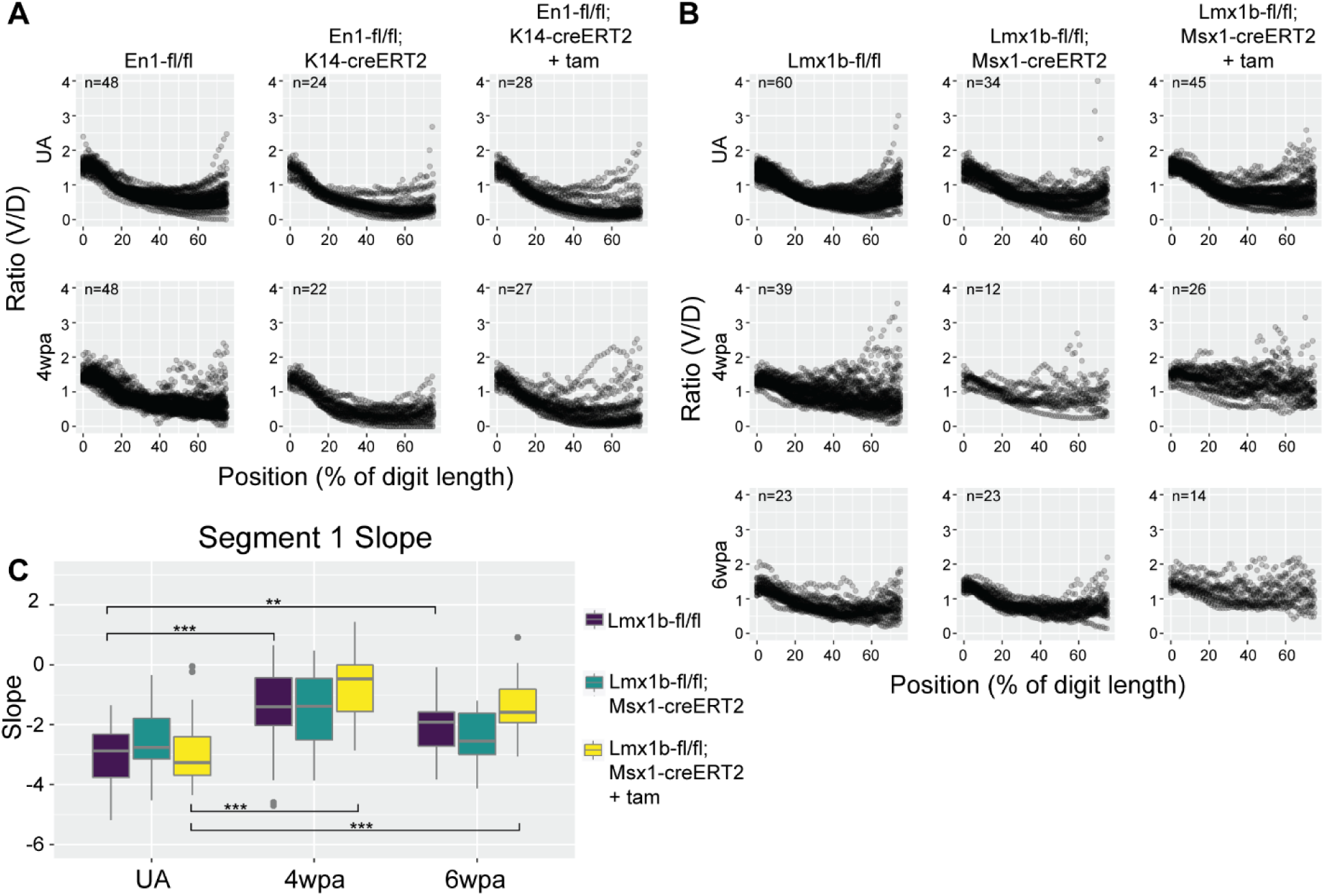
DV symmetry in individual digit tip bones. (A) Raw ventral/dorsal volume ratio measurements for En1-fl/fl, En1-fl/fl; K14-creERT2, and En1-fl/fl; K14-CreERT2 + tam unamputated (top) and 4wpa (bottom) digits. (B) Raw ventral/dorsal volume measurements for Lmx1b-fl/fl, Lmx1b-fl/fl; Msx-creERT2, and Lmx1b-fl/fl; Msx-CreERT2 + tam unamputated (top), 4wpa (middle), and 6wpa (bottom) digits. n denotes the number of individual digits in each group. (C) Box plot showing the mean slope predictions of segment 1 for the Lmx1b cohort. * p < 0.05, ** p < 0.01, *** p < 0.001.

